# Transcription factor occupancy is affected by genetic variation in the white adipose tissue of C57BL6/j and 129S1/SvImJ mice

**DOI:** 10.1101/2025.05.08.652810

**Authors:** Juho Mononen, Mari Taipale, Marjo Malinen, Anna-Liisa Levonen, Anna-Kaisa Ruotsalainen, Luke Norton, Sami Heikkinen

## Abstract

Most of the disease associated genetic variants identified in genome wide association studies have been mapped to the non-coding regions of the genome. One of the leading mechanisms by which these variants are thought to affect disease susceptibility is by altering transcription factor (TF) binding. Even though inbred mouse strains have been commonly used to investigate polygenic diseases, less is known on how their genetic differences translate to the level of gene regulation and chromatin landscape. Here, we investigated how genetic variation affects chromatin accessibility, a commonly used proxy for TF binding, and how this relates to gene regulation in the epididymal white adipose tissue (eWAT) of C57BL/6J and 129S1/SvImJ mice fed either chow or high-fat diet. We show that differences in chromatin accessibility are almost exclusively strain-specific and driven by genetic variation. In addition, we integrate ATAC-seq (chromatin accessibility) and H3K27ac ChIP-seq (active regulatory regions) data to show that tissue-specific TFs are commonly found in the active regulatory regions hosting TF binding motif altering variants in eWAT. By incorporating footprint analysis, we also show that TF occupancy is consistent with TF binding motif scores at the genetically altered loci. In addition, we validate these findings by extending the footprint analysis to ATAC-seq and H3K27ac ChIP-seq data obtained from the liver. Also, we employ RNA-seq to show that differentially expressed genes are collocated with differentially accessible regions hosting genetic variants. Overall, our findings highlight the connection between differential chromatin accessibility, TF binding and genetic variation and their role in the gene regulation across metabolically central tissues of a mouse model for polygenic obesity.

**Author summary:** Most of the variants associated with human polygenic disease are outside the protein coding regions of the genome. In this study, we took benefit of the known genetic differences of two inbred mouse strains to investigate the impact of the non-coding genetic variation on the chromatin landscape and the regulation of gene expression in the adipose tissue. Using several complementary sequencing and bioinformatic methods, we show that chromatin accessibility at genetically different transcription factor (TF) binding sites is associated with the strength of the altered TF binding motifs. We also identify the key TFs binding to the regions of genetically determined chromatin accessibility. We further show how the genetic effects propagate to the level of gene expression. Together, these findings give valuable insight into the mechanisms by which genetic variation affects gene regulation and provide future targets for research.

## Introduction

Inbred mouse models have been used to study the mechanisms behind polygenic diseases such as multifactorial obesity [1]. However, most of the research has focused on identifying the key regulatory regions and the underlying biology for these conditions whereas the more subtle effects of genetic variation have not been thoroughly studied [1–3]. As most of the single nucleotide polymorphisms (SNPs) found in genome-wide association studies on e.g. polygenic obesity are located within the non-coding genome, future research would benefit from understanding how these variants effect the gene regulatory landscape [4]. One of the proposed mechanisms by which these regulatory SNPs (rSNPs) work is that they alter transcription factor (TF) binding [5,6]. Most recently, several studies have demonstrated this phenomenon, but picture is far from complete e.g. on the predominantly affected TFs and their target genes [5–7].

Previously, differential chromatin accessibility has been shown to be affected by genetics but not diet [2,6]. Assay for Transposase-Accessible Chromatin sequencing (ATAC-seq) has been used widely to assess the effects of genetic variation to regulatory landscape as it can be used to detect candidate TF binding sites and has been shown to correlate well with nearby gene expression [6,8–10]. Distinct classes of TFs relate differently to the regulation of chromatin accessibility. For example, pioneer factors are a group of TFs which can initiate the opening of closed chromatin regions and thus activate gene expression by allowing other TFs, such as settlers and migrants, to bind chromatin [11]. As TFs bind in a sequence-specific manner, genetic variation at the binding sites is known to affect TF binding and, in the case of pioneer factor binding, chromatin accessibility [5,6,12]. Even though pioneer factors are known to function in cell type specific manner, little is known about how genetic variation affects their function and how this propagates to the tissue level. Recent studies have focused more on the analysis of gene expression data when estimating the effects of genetic variation on the events contributing to the differentiation of cell types and thus additional studies to identify cell type specific pioneer factors and how their function is affected by genetic variation are needed [13,14].

As shown previously by us and others, differentially accessible regions (DARs) are often enriched for genetic variation and disrupted TF binding motifs [6,8,9,15]. However, looking purely at altered TF binding site motifs in DARs has provided poor accuracy for characterizing rSNPs with roughly ∼20% of altered motifs disrupting TF binding [5,6]. For better accuracy, additional methods, such as ChIP-seq, are needed, making the identification of novel interactions cumbersome [6,16]. To study the complex landscape of polygenic diseases where multiple different TFs can be affected, more comprehensive methods for identifying candidate rSNPs must be explored. Footprint analysis is an *in silico* method that identifies TF occupancy sites from accessibility sequencing assays such as ATAC-seq [17]. A recent study by Viestra *et al.* highlighted the usefulness of footprinting in identifying candidate rSNPs by showing that TF footprints were enriched for genetic variation compared to the rest of the non-coding genome [18]. However, footprinting has not been tested in identifying regulatory variants in inbred mice.

In this research we study the relationship of genetic variation and the regulatory landscape in epididymal white adipose tissue (eWAT) of C57BL/6J (B6), a widely used mouse model in diet-induced obesity research, comparing it to 129S1/SvImJ (129) mice [1]. We show that in eWAT the genetic effects on chromatin accessibility are highly strain-specific and independent of diet-induced effects. We identify active regulatory regions from ATAC-seq and H3K27 acetylation (H3K27ac) ChIP-seq data using an integrative approach which has been previously shown by us and others to increase the accuracy *in silico* TF binding site identification [6,19]. Moreover, we use RNA-seq to show that DARs are located near, and correlate with, differentially expressed genes (DEGs) especially at the promoters. Using footprint analysis of ATAC-seq data, we further show that genetically determined chromatin accessibility co-locates with altered TF binding motifs and that ∼75% of the motif scores are altered conjointly with the local accessibility. For additional validation, we expand the footprint analysis to our previously published dataset on the genetically driven regulatory landscape in the liver [6]. These findings also present a compelling case for future studies focusing on genetic variation and pioneer factor driven cell type differentiation to better characterize the cell type specific pioneering TFs and their role in the regulatory cascades of gene expression.

## Results

### Differences in chromatin accessibility are evident between the strains but not induced by the diet

We discovered 98,214 nucleosome-free regions (NFRs) across the samples (5-6 per diet and strain). Preliminary analysis suggested a possible confounding within the sample set (**Fig S1A**). Enrichment analysis for the DARs (FDR<0.01 and fold change > 50%) which were more accessible in Cluster 2 showed enrichment for spermatozoa and epididymal cell specific genes when cross-referenced with a single cell reference for B6 eWAT (**Fig S1B**) [20]. Therefore, we performed our main DAR analysis using csaw by batch-correcting for epididymis cluster membership which successfully eliminated the bias (**Figs S1C-D**). Regardless, almost no diet-induced changes in chromatin accessibility were observed in either strain whereas hundreds of inter-strain DARs were detected (**Fig 1A**). About one third of the 4,105 DARs were common in the two diet comparisons (**Fig 1B**). Due to the scarcity of diet-induced changes, in subsequent analysis we focused on the inter-strain DARs, classifying each as ‘Chow-DAR’ (DAR only on chow), ‘HFD-DAR’ (DAR only on HFD), or ‘Common-DAR’ (DAR on both diets), and dividing the classes further, when applicable, to up and down DARs (**Fig 1C**). We also performed enrichment analysis for the DAR-classes to neighbouring genes using GREAT (**Fig 1D**). Both Chow-and Common-DARs were enriched, either specifically or together, for GO biological processes related to actin filaments, such as “Actin filament bundle assembly”. This is interesting because *e.g.* actin depolymerization has been characterized as a key step in adipocyte differentiation [21]. On the other hand, the HFD-DARs were specifically enriched to processes of lipid metabolism and Bone Morphogenetic Protein (BMP) signalling, which is known to be connected with adipose tissue function and adipogenesis [22].

**Fig 1.**
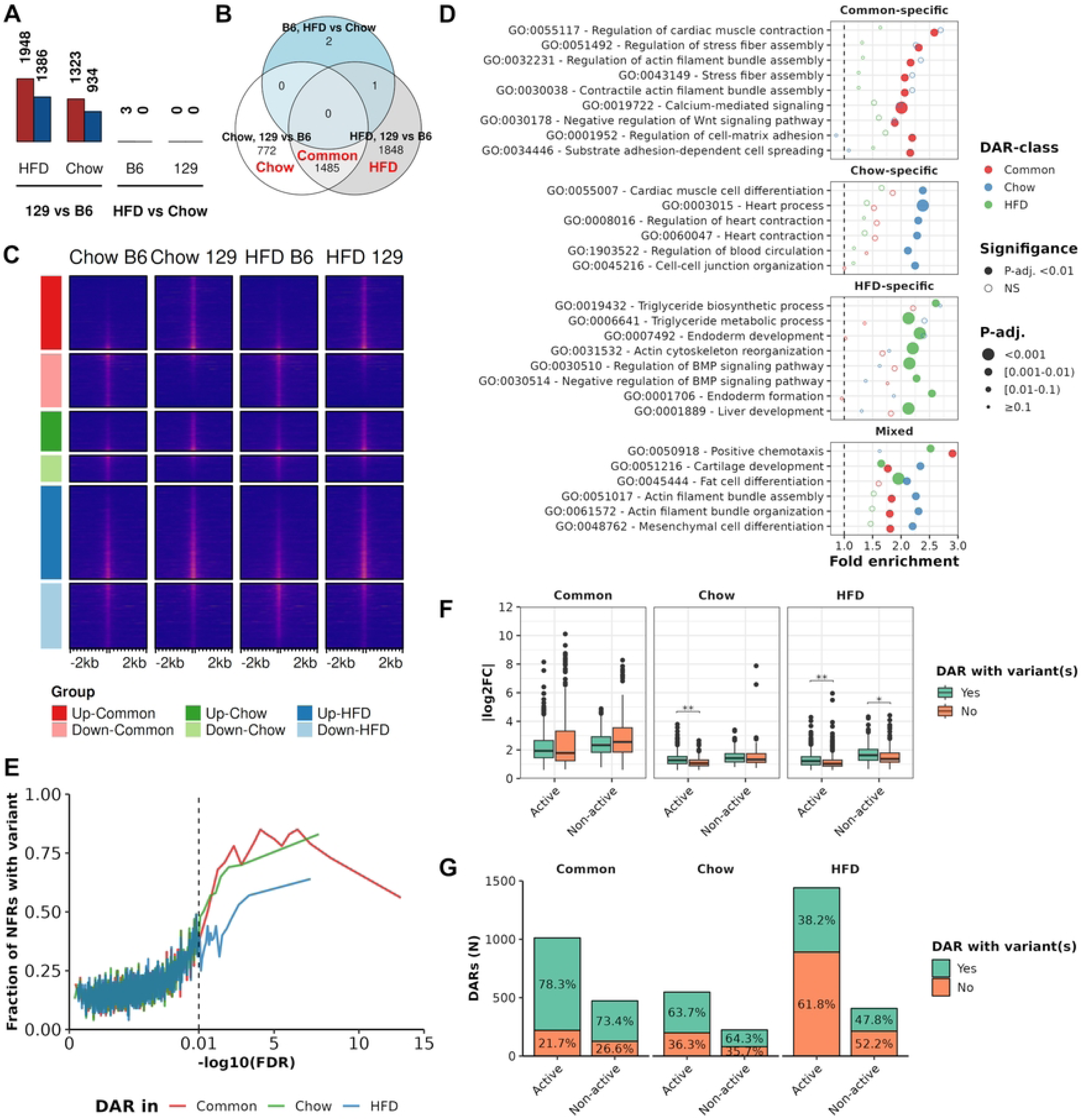
Differences in chromatin accessibility are almost exclusive between the strains and enriched for genetic variation. **A)** Counts of DARs observed in different comparisons. **B)** Venn diagram of DAR counts in comparisons with any DARs. DAR-classes^1^ labelled with red. **C)** ATAC-seq signal enrichment in different groups across the DAR-classes^1^. **D)** Results from GREAT analysis of DAR-classes^1^. Top 10 pathways (with hypergeometric test FDR<0.01) were selected from each DAR-class and results were plotted for all classes. **E)** The fraction of ATAC-seq peaks containing at least one genetic variant (y-axis) in windows of 100 NFRs ordered by the significance of difference within diet comparisons (x-axis). FDR for Common-DARs is for the more statistically significant comparison (129 vs B6 on either chow or HFD). **F)** Boxplot of absolute log2 fold changes in different DAR-classes comparing between DARs with and without variants (Wilcoxon test P-value, **=<0.01 and *=<0.05). **G)** Counts of DARs (y-axis) and percentages of DARs with variants across (x-axis). ^1^Common=”DAR in both diet comparisons”, HFD=”DAR in HFD comparison”, Chow=”DAR in chow comparison”

### Largest differences in chromatin accessibility are driven by genetic variation

Differential accessibility displayed high concordance with genetic variation as higher significance DARs were more likely to overlap genetic variants (**Fig 1E**). Interestingly, the DARs observed only on one diet had somewhat different characteristics than the Common-DARs. For example, Common-DARs had generally higher absolute fold-changes than the Chow- or HFD-DARs (**Fig 1F**). We categorized those NFRs as active that were surrounded by H3K27 occupancy, and the rest as non-active [6]. Separating DARs into active and non-active provided only small differences, with active Chow- and HFD-DARs displaying higher fold changes than non-active in the presence of variants while similar effect was not observed with Common-DARs (**Fig 1F**). However, Common-DARs were more enriched for genetic variants than either the Chow- or HFD-DARs (**Fig 1G**). This difference was especially clear for HFD DARs, 40.3% of which overlapped with genetic variants, compared to the 76.3% of Common-DARs. Notably, between the diet categories, active and non-active DARs had roughly similar proportions of overlapping SNPs.

### Differential accessibility propagates to local chromatin landscape

Next, we wanted to investigate how the differential accessibility relates to the chromatin landscape. As expected, NFRs classified as active displayed higher flanking H3K27ac signal (**Fig 2A**). The distributions of active and non-active categories were similar in all DAR-classes regardless of the presence or absence of variants at the NFR (**Fig 2B**). In all DAR classes, including non-DARs (non-DAR in all comparisons), and at all types of genomic features, active NFRs were more accessible than non-active NFRs (**Figs S2A and S2B**). Interestingly, active DARs with and without variants were more localized to distal regulatory regions than were non-DARs (**Fig 2C**), even if in general the chromatin was most accessible at the promoter regions of expressed genes (**Figure S2B**). DARs also often correlated positively with their nearest DARs (**Fig 2D**). Interestingly, however, DARs with variants were less correlated with their nearby DARs with variants compared to DARs without variants, and positive correlation was most frequent, and occurred over greater distances, between DARs that both are without variants (**Fig 2D**). Collectively these findings suggest that the disruption of chromatin accessibility caused by genetic variation spreads up to 250kb into local chromatin landscape.

**Fig 2.**
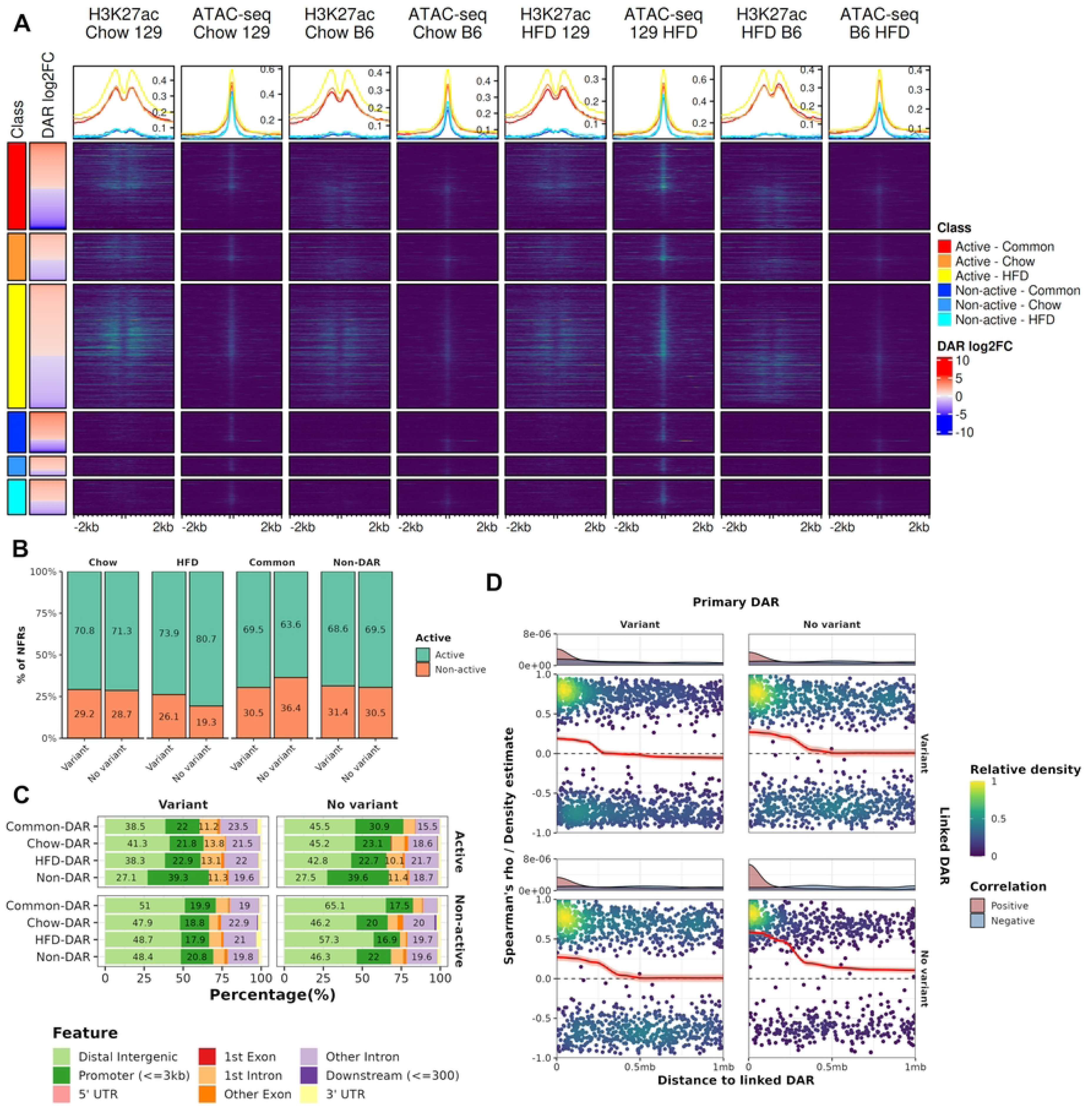
Differential chromatin accessibility locates in distal regulatory regions and spans to nearby NFRs. **A)** Enrichment of ATAC-seq and H3K27ac ChIP-seq signal at DAR-classes^1^ across different groups. **B)** Percentages of NFRs with and without variants in active and non-active NFRs. **C)** Genomic distribution of NFRs from different classes. Percentages>10% labelled. **D)** Scatter plot for Spearman correlation coefficients for DARs (Primary DARs) and their neighbours (Secondary DARs) with and without variants. Distribution of values presented with locally estimated scatterplot smoothing (red line, ±SE), and the densities of negatively and positively correlated secondary DARs are shown above each scatter plot. ^1^Common=”DAR in both diet comparisons”, HFD=”DAR in HFD comparison”, Chow=”DAR in chow comparison”, Non-DAR=“Non-DAR in both comparisons”

### Differential gene expression is more pronounced between the strains than diets

To enable linking the differences in chromatin accessibility to gene expression, we generated eWAT RNA-seq data for the same mice as for ATAC-seq. Similar to ATAC-seq, preliminary analysis of cluster-wise differentially expressed genes (DEGs, FDR<0.05, **Table S1**) suggested confounding of epididymis origin (**Figs S3A and S3B**). After successful batch correction (**Figs S3A and S3C**), the differences between the strains were again the most numerous and with higher effect sizes compared to differences between diets (**Fig 3A**). For downstream analyses, we categorized genes to four classes; Non-DEG (non-DEG in all comparisons), Diet-DEG (DEG only in at least one diet comparison), Diet+strain-DEG (DEG in at least one diet and one strain comparison) and Strain-DEG (DEG in both strain comparisons). Interestingly, strain-DEGs were most enriched to immune system related pathways whereas Diet+Strain DEGs and Diet DEGs were mostly enriched to pathways related to metabolism (**Fig 3B**).

**Fig 3.**
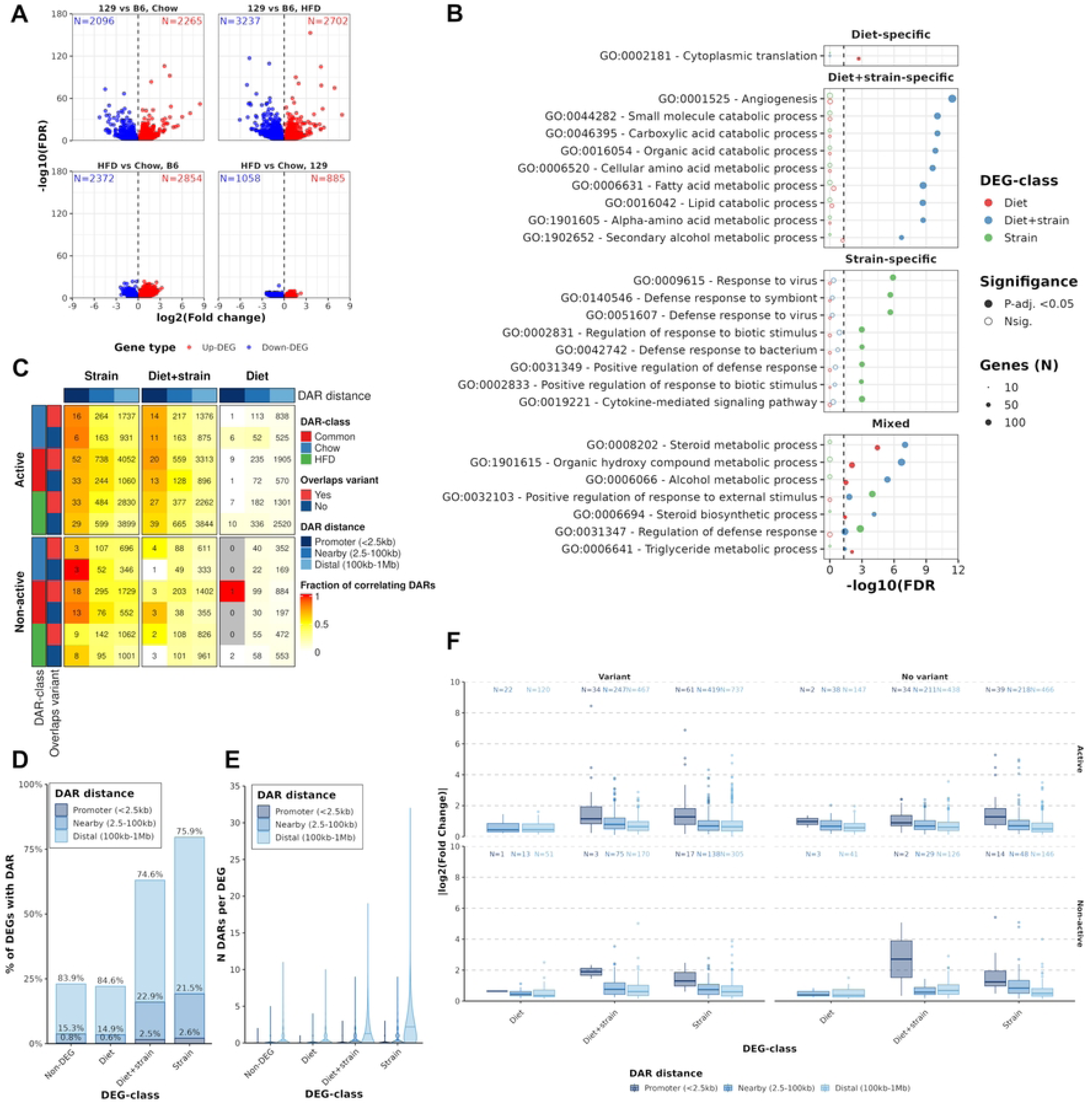
Strain-specific gene expression is connected with genetically determined chromatin accessibility. **A)** Volcano plots of differentially expressed genes by comparison. **B)** GO-term over representation analysis results for DEG-classes using expressed genes as the background. Top 10 pathways (with FDR<0.05) were selected from each DEG-class^1^. **C)** Heatmap displaying fractions of DARs^2^ correlating (Spearman’s Rho>0 and P-value<0.05) with DEGs in different windows from the active TSS of the DEG*. Total count of DARs annotated with numbers. **D)** Fraction of expressed genes with DAR (of any class) in different windows around their active TSS. Percentages displayed are for fractions per DEG-class^1^. **E)** Counts of correlating DARs (of any class) in the neighbourhood of genes by DEG-class^1^. **F)** Absolute log2 fold-changes of DEGs^1^ with nearest DAR (of any class) in different windows around active TSS. ^1^Non-DEG=non-DEG in all comparisons, Diet=DEG only in diet comparison, Diet+strain=DEG in both diet and strain comparisons, Strain DEGs=DEG in both strain comparisons. ^2^Common=”DAR in both diet comparisons”, HFD=”DAR in HFD comparison”, Chow=”DAR in chow comparison”

### Differentially expressed genes have multiple correlating accessible regions in their neighbourhoods

When DARs were co-located with DEGs, the strongest correlation between chromatin accessibility and gene expression was observed at the promoters for both Strain- and Diet+Strain-DEGs, where ∼80% of DARs correlated positively with DEGs (Spearman’s correlation, P-value<0.05, Rho>0) (**Fig 3C**). Active DARs tended to correlate more often with nearby DEGs than non-active DARs (**Fig 3C**). Strain- and Diet+strain-DEGs, compared to Non- and Diet-DEGs, much more often had correlating DARs in their neighbourhoods, especially at shorter distances (<100kb) (**Fig 3D**), and there also were more of them per DEG (**Fig 3E**). Interestingly, Diet-DEGs appear independent of DARs since the percentages of them with DARs closely resembled those of Non-DEGs in all distance categories (**Fig 3D**). We also wanted to assess how NFRs are generally found near DEGs regardless of correlation. Linking genes to their nearest NFR provided only minimal differences between the groups (**Fig S3D**). However, both Strain- and Diet+strain-DEGs with variant-overlapping DAR at their promoter displayed more extreme fold changes than Diet-DEGs, or corresponding categories where the DAR had no variant, further highlighting the likelihood of regulatory variation (**Fig 3F**). Since the distance to the nearest NFRs tended to be longer for genes with lower expression, the incomplete overlap of NFRs and promoters of expressed genes is likely to be explained by the suboptimal sensitivities of both ATAC-seq and H3K27ac ChIP-seq at the promoters of low expressed genes (**Figs S3E and S3F**). These findings confirm the strong link between chromatin accessibility and gene expression and suggest that regulatory actions from outside the proximal promoter are a major contributor to genetically determined gene expression.

### Motif analysis connects chromatin accessibility to cell type specific transcription factors

After confirming the strong link between chromatin accessibility and gene expression, we looked for key factors affecting the gene regulatory landscape. Active NFRs were enriched for motifs of known WAT-associated TFs like CEBPs, AR and NR3C1 (also called GR), whereas non-active NFRs displayed high enrichment of the CTCF binding motif, highlighting their potential roles in establishing the chromatin conformation (**Fig S4A**) [23,24]. Among DARs, non-active DARs of all classes were similarly more enriched for CTCF binding motifs than active DARs (**Fig 4A**). In contrast, binding motifs for SPIC, IRF4 and ETS2 were enriched in active Common- and HFD-DARs over all non-active and Chow classes. However, both non-active and active HFD-DARs were enriched for AR and STAT5A binding motifs over other classes. Suspecting that the differences in cell type composition might contribute to the differences between the DAR classes, we performed a chipenrich analysis of DARs and cell type specific genes. Indeed, the HFD-DARs were the only class of DARs to be enriched to cell type specific genes, namely adipocytes and immune cells (**Fig S4B**). Combined with the observation that the HFD-DARs had genetic variants less often than the other two DAR classes (**Fig 1D**), this suggests that the HFD-specific set of DARs is enriched for cell-type specific sites that were detected at least partly due to the increased relative amount of adipocytes and immune cells in the obese B6 eWAT. In summary, DARs with variants might be driving the accessibility of DARs without variants, but accurate interpretation of these observations is hindered by the presence of cell type specific DARs.

**Fig 4.**
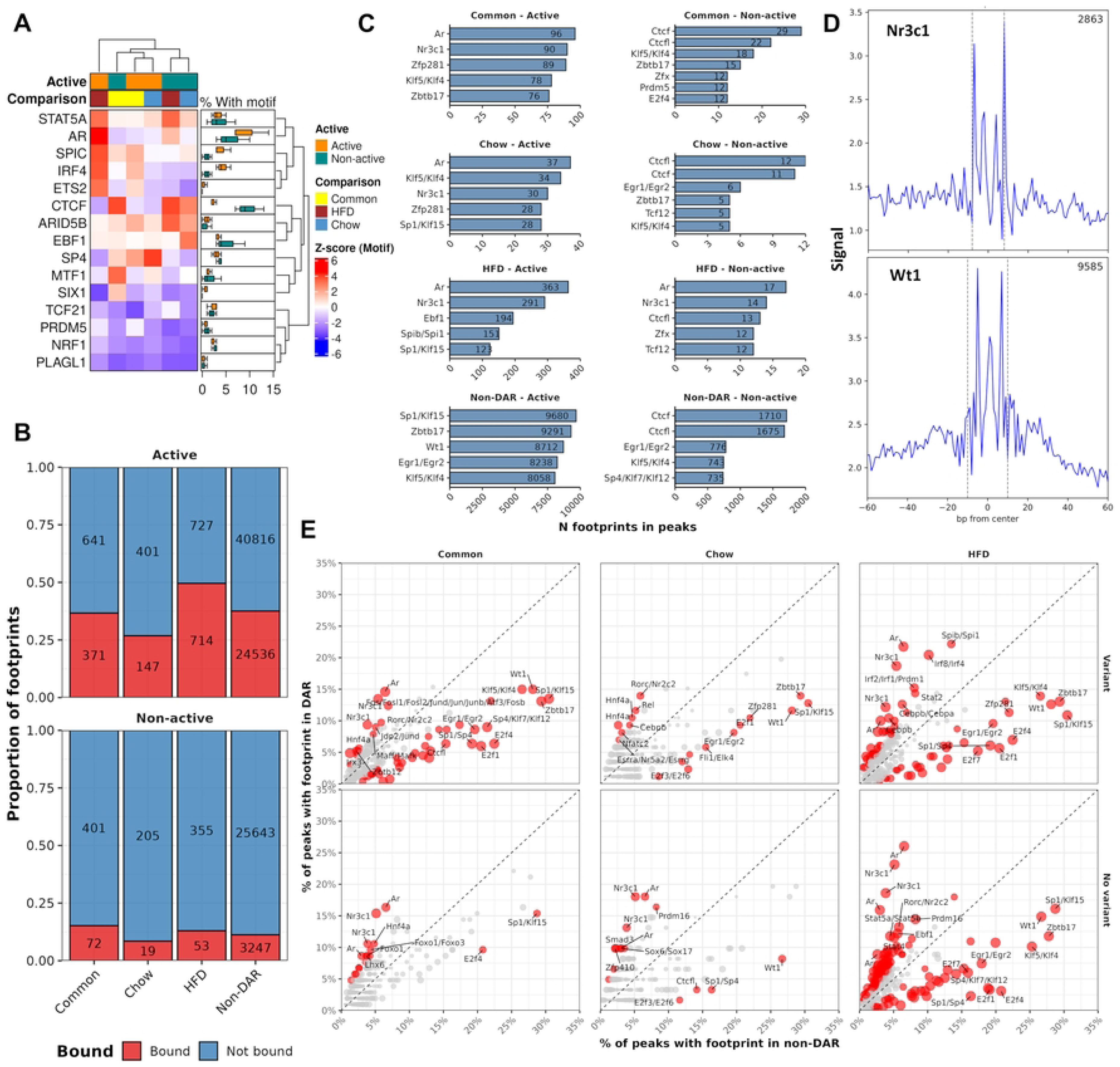
Accessible chromatin overlaps with known adipose tissue transcription factor footprints. **A)** Heatmap of differential motif enrichment in DAR-classes^1^. Boxplot on the right summarizes the percentages of active and non-active NFRs with motifs across the DAR-classes. **B)** Fractions of NFRs^1^ that overlap with at least one footprint in eWAT. **C)** Counts for top 5 most common footprints in active and non-active NFRs^1^ in eWAT. **D)** Footprint aggregates for Nr3c1 and Wt1 across all detected footprints in all groups in the NFRs^1^. **E)** Scatter plot of footprint occupancy in DARs^1^ vs non-DAR. Size of points is determined by Chi-squared P-value; red, P-value<0.05. Top 10 TFs with highest difference in occupancy fraction between DAR-class and non-DAR are labeled (P-value < 0.05). ^1^Common=”DAR in both diet comparisons”, HFD=”DAR in HFD comparison”, Chow=”DAR in chow comparison”, Non-DAR=“Non-DAR in all comparisons”.

### Footprint analysis reveals binding sites for tissue-relevant factors

To better understand the results of motif enrichment analysis and to address TF binding more reliably at NFRs, we performed TF footprint analysis using TOBIAS [17]. Footprints identified in the different groups were largely overlapping (**Fig S5A**). Interestingly, the largest individual groups of footprints were observed with the two HFD groups, which is not surprising since HFD DARs were the largest DAR group. As expected, active NFRs of all classes hosted footprints more frequently than non-active NFRs (**Fig 4B**), further highlighting the benefit of separating NFRs based on the flanking H3K27ac signal. Additionally, among the most common footprints in non-active NFRs were footprints for CTCF and its’ paralog CTCFL (**Fig 4C**). In contrast, active NFRs hosted footprints for several well-known TFs relevant in WAT such as NR3C1 (*i.e.* GR, lipolysis and immune response), AR (adipocyte proliferation and metabolism) and WT1 (lipid metabolism) (**Figs 4C and 4D**) [24,25]. Interestingly, AR was also present in many non-active HFD-DARs. We also tested whether a given footprint was more common in active DARs compared to active non-DARs. We limited this analysis to those NFRs that had any footprints because the footprint scores, and thus binding classification, depends heavily on the ATAC signal at the NFR, and because substantial differences between the NFR classes were observed (**Fig S5B**). Quite strikingly, across all DAR sets the results highlighted more frequent binding of AR and NR3C1 in DARs compared to non-DARs (**Fig 4E**). Interestingly, NR3C1 is classified as a pioneer factor and has been known to partake in the regulation of AR activity in human adipocytes [24]. However, the motifs for both AR and GR are similar which makes their accurate profiling by footprinting difficult (**Fig S6A**). Similarity of motifs for other conjointly enriched TFs was also observed (**Fig S6A**). On the other hand, WT1, which was more frequently bound in non-DARs, is an important regulator for the normal function of WAT, disruption of which can lead to severe developmental deficits [25,26]. As expected, higher proportion of the TFs enriched in DARs were either Strain- or Diet+strain-DEGs compared to those enriched in non-DARs (**Fig S6B**). Interestingly, of the DAR-enriched eWAT TFs, especially those which were enriched in DARs without variants presented themselves as cell type specific markers in a single cell reference more often than the other groups (**Fig S6C**).

### The footprint patterns of eWAT are replicated in the liver

To expand the analysis to a larger set of TFs and metabolic functions, we replicated the footprint analysis on our recently published liver dataset [6]. Similar to eWAT, also in the liver the footprints largely overlapped between the groups (**Fig S7A**). As expected, because the Chow-DARs was the largest DAR group, chow groups had more unique footprints than respective HFD groups (**Fig S7B**). The active NFRs hosted more footprints, the most frequent being the known hepatic regulators such as HNF4a and the ETS TF family members (**Figs S7B and S7C**) [27,28]. As expected, CTCFL was the most frequent footprint among the non-active NFR-classes. The enrichment analysis concurred with the frequencies, as *e.g.* HNF4a was highly overrepresented in Chow-DARs (**Fig S7D**). Interestingly, the liver results replicated the enrichment of SP1 and WT1 footprints in non-DARs also seen in eWAT (**Fig S7D**). However, the motifs for SP1, a known transcriptional regulator in both eWAT and liver, are similar to WT1 (**Fig S6A**) [29,30]. Similar patterns seen in both tissues further strengthens the view that genetically determined chromatin accessibility is mediated by a certain set of TFs displaying pioneer activity in the tissue.

### Genetic variation is driving differences in transcription factor binding at differentially accessible regions both in eWAT and liver

Next, we investigated the impact of genetic variation in the differential TF binding and chromatin accessibility. Similar to our previous findings in liver, altered motifs in the direction of their overlapping Common-DARs were enriched for the motifs of key factors mediating enhancer action *e.g.* CEBPB and CEBPA (**Fig 5A**) [6,31]. There was no significant enrichment for any altered TF motif in the Chow- or HFD-DAR classes (data not shown). In active NFRs, we identified 1,480 and 573 footprints for eWAT and liver, respectively, where the motif allelic score and the corresponding footprint score had the same direction of effect (altered motif corresponding footprint, AC-footprint), (**Figs 5B and 5C, Table S2**). Among the DAR classes, the two scores concurred for most motif-footprint pairs except for the HFD-DARs in liver. When the distribution of footprint score differences between the strains were evaluated over the bins of motif score differences, for the non-DARs the footprint score differences showed no concurrency with the motif score differences even in the most extreme bins neither in eWAT nor the liver, whereas for the DAR-classes the concurrency between the two score differences agreed with the more simpler frequency-based analysis (**Figs 5D and S8A**). Notably, however, the magnitudes of the footprint score differences did not increase or decrease towards the more extreme bins of motif score differences. As highlighted before, footprint scores have a strong positive correlation with the ATAC-seq signal and are also affected by cell type specific binding effects, which could influence the detection of genetically determined binding using footprint analysis in non-DARs (**Figs S5B and S6C**).

**Fig 5.**
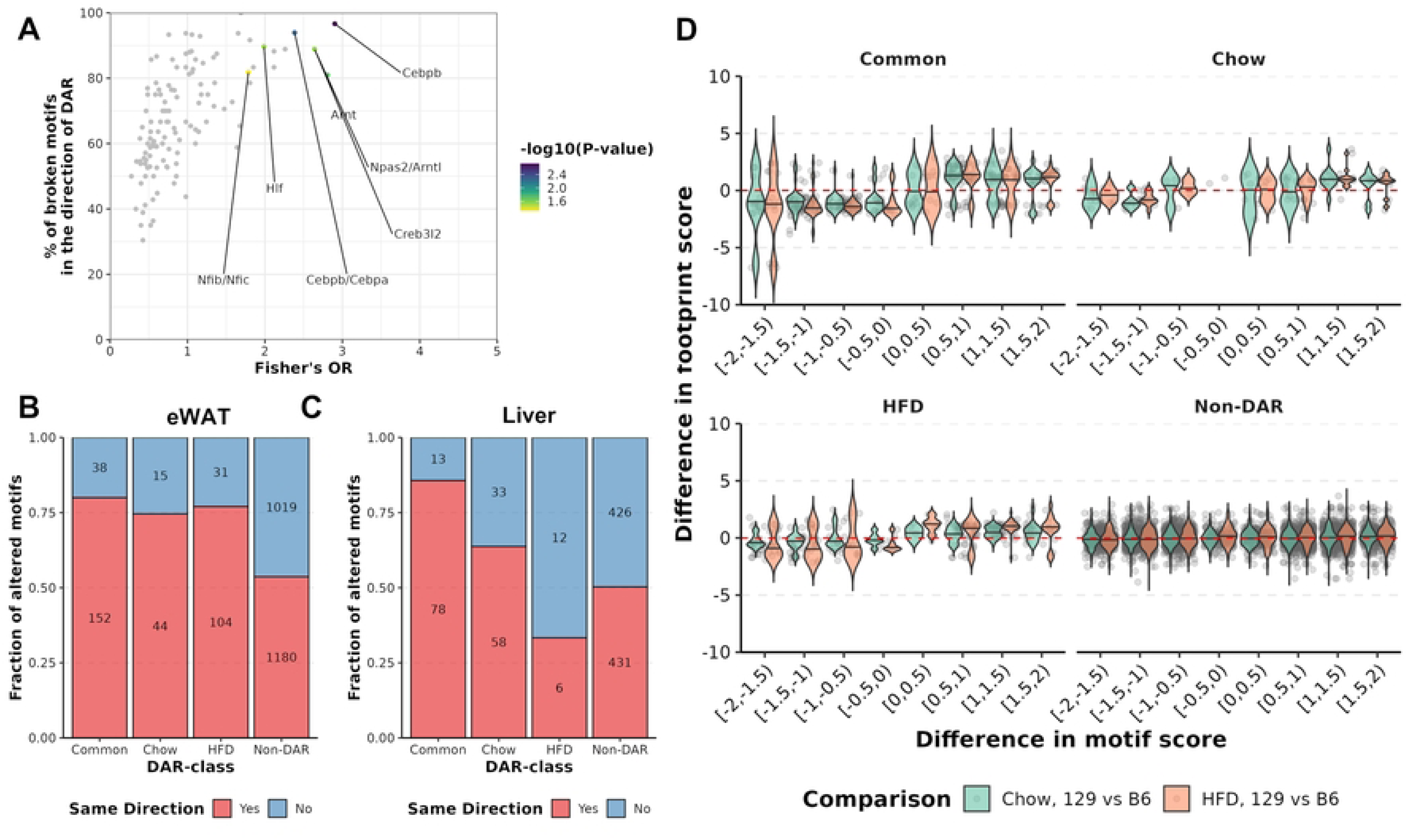
Footprint scores are predictive of motif-altering genetic variation in DARs. **A)** Comparison of variant-altered *(i.e.* broken) TF binding motifs between Common-DARs and non-DARs using Fisher’s test. TFs with *P*-value < 0.01 are coloured, and further those with >80% of altered motifs with collateral motif score to DAR log2FC and *P*-value <0.01 are labelled. Fractions of altered motif overlapping footprints with motif score in the direction of footprint score in NFR-classes^1^ for **B)** eWAT and **C)** liver. Counts of footprints are given within the bars. **D)** Differences in eWAT footprint scores (Y-axis) across Motif allelic score difference (129 vs B6) categories (X-axis). ^1^Common=”DAR in both diet comparisons”, HFD=”DAR in HFD comparison”, Chow=”DAR in chow comparison”, Non-DAR=“Non-DAR in both comparisons”.

Finally, we focused on the identification of pioneering factors among the altered motifs overlapping with footprints in DARs, to help us understand whether the genetic variation at chromatin remodelling TF binding sites contributes to differential accessibility. The most common AC-footprints in eWAT DARs were those for *e.g.* JDP2/JUND and ETS-family factors which have been suggested, or are known, to have pioneer activity (**Fig S8B**) [32,33]. In the liver the results were quite similar such that the ETS-family factors were commonly present among the most frequent AC-footprints (**Fig 7C**). Unfortunately, due to the low number AC-footprints per TF (**Fig S8D**), inferring the over-representation of pioneer factors in the footprints of DARs provided only minimal results. Only one TF in eWAT (STAT6, Common-DAR vs Non-DAR Fisher’s P-value=0.042) was significantly enriched. However, many of the TFs close to the selected cut-off for significance (P-value<0.05) were mostly concordant by footprint and motif score (**Figs S8E and S8F**). To summarize, the altered motifs for TFs known to exhibit pioneer activity could be found in the DARs and non-DARs. In addition, the presence of motif-altering variants at footprint was predictive of differential binding at DARs.

## Discussion

In this study we expand on our previous analysis in liver and show how differential chromatin accessibility links to genetic variation and gene expression also in eWAT using a combination of ATAC-seq, RNA-seq and H3K27ac ChIP-seq data [6]. We also inspect how *in silico* inferred binding sites from TF footprinting can be used to detect genetically determined TF binding.

Similar to our findings in liver, differential accessibility between 129 and B6 mice on chow and HFD was only observed between the strains in eWAT. Regions displaying differential accessibility, especially the most significant ones, were also co-localized with genetic variation. However, unlike in liver, in eWAT we did see pronounced effects of cell type composition affecting both ATAC-seq and RNA-seq samples. Adipose tissue samples have been shown to have highly variable cell type proportions due to tissue heterogeneity, adjusting for which generates more accurate downstream results [34]. Indeed, we identified a subset of samples with significant epididymis-related cell enrichment and employed batch correction for more accurate downstream analysis. Due to the lower number of H3K27ac samples we did not incorporate batch correction there. However, regardless of the batch correction, some signs of differences in cell type composition seem to remain *e.g.* because AR and GR footprints were somewhat surprisingly observed also in non-active HFD-DARs, suggesting that the effects of cell type composition propagate also to the H3K27ac signal.

Interestingly, while a strong correlation between DARs without variant and their nearest DAR with variant was observed, DARs without variants presented stronger correlation with other DARs without variants. Given that DARs without variants were most prominent in HFD-DARs which were enriched near adipocyte and immune cell specific genes, it is possible that the stronger correlation is due to differences in the cell type composition as B6 mice have a more pronounced adipose tissue expansion on HFD [1]. In addition, although the existence of multi-NFR chromatin accessibility QTLs (caQTL) has been demonstrated, they are rare when compared to genetically determined single NFRs [15]. In the study by Gate et al. which focused only on CD4+ T-cells, and was thus void off any cell type composition effects, co-accessible peaks only comprised 2% of the total observed NFRs [8]. In addition, it could be possible that the lower correlation between DARs with variant and other DARs is explained by the disruption in the local chromatin landscape mediated by altered TF binding, or that the neighbouring variants not overlapping with DAR could affect chromatin accessibility [15].

DARs of all classes were predominantly distal to DEGs, but largest effects on gene expression were seen when a DAR with variant was localized at the promoter of a gene. Although DARs were numerous in the neighbourhoods of Strain- and Diet+strain-DEGs, outside of the promoter the fraction of correlating DARs quickly dropped, highlighting that with bulk transcriptomics data the distal NFR-to-gene interactions are hard to interpret. In addition, some sensitivity issues were noticed with the promoter accessibility of lower expressed genes, which may have excluded some of their TSS’s from promoter-based analyses. Single nucleus ATAC-seq has been shown to solve some of the aforementioned issues on cell type specific chromatin accessibility when detecting genetically determined chromatin co-accessibility [35]. However, based on previous studies on both caQTLs and integrated analysis of gene expression and caQTLs, the strongest correlating changes have been observed at short distances, which suggests that the analysis of the proximal connections between chromatin accessibility and gene expression are fairly reliable [8,9,15,35].

Interesting observations were made from the enrichment of footprints between different NFR-classes. For example, DARs in both tissues were enriched for TFs that are known to partake in the differentiation of tissue-specific cells. In eWAT, the highest enrichment in DARs (compared to non-DARs) was observed for GR and AR. The binding motifs for these two TFs are similar, making it difficult to distinguish the actual effector [36]. Interestingly, however, GR is known to regulate AR expression in WAT and is an important factor in adipogenesis and it is also known to present pioneer activity, which could hint that it is GR rather than AR that regulates accessibility in eWAT [24,37]. In liver, the most strikingly enriched TF was HNF4α which is a key factor in regulating hepatic progenitor cell formation [38]. Interestingly, neither GR or HNF4α dysfunction has been shown to have severe phenotypic consequences in their respective tissues [37,39]. In contrast, WT1, which was enriched in non-DARs in both tissues, is an important lineage-defining TF [25,40]. In healthy liver, WT1 is expressed only in the fetal stage where it is an important factor for mesenchymal lineage determination [40]. In WAT, WT1 is necessary for maintaining adipose identity and it is an important factor regulating fat cell development, the dysregulation of which leads to severe developmental deficits [25,41]. This observation could suggest that differential accessibility is driven by more dispensable TFs that could affect cell type composition while alteration in the binding sites of TFs that are required for normal development might not present differential accessibility in adult liver due to more severe consequences. However, it is important to note that the expression of WT1 in adult liver is only observed after liver damage and fibrosis which can affect the conclusions made here [42]. Additionally, WT1 sharing a similar motif with another lineage determining factor SP1 could make interpreting these results difficult [29,30]. We observed several cell type specifically expressed TFs among the enriched TF footprints in DARs not overlapping with genetic variation, suggesting that DARs with variants are less cell type specific, while DARs without variants are more likely to be reflected in cell type composition differences. As TF expression has been shown to be affected by genetic factors and result in differences in cell type differentiation, additional studies on both *cis*- and *trans-*acting variants on cell type specific pioneer TF function are needed [13,14]. However, the overall number of enriched TFs was quite low for DARs with variants which limits the reliability of this interpretation. In summary, TF binding sites for tissue specific TFs partaking in both the regulation and cell type differentiation were often found in DARs, highlighting the benefits of incorporating footprint analysis in the detection of accessibility dependent TFs.

Footprint analysis gave us more insight on which TFs bind at the DARs and non-DARs. However, while the enrichment of altered motifs in DARs vs non-DARs was observed to have similar patterns than in our previous work with liver, enrichment of AC-footprints in DARs did not provide much additional insight [6]. Although we saw CEBP-family TF binding motifs enriched in the altered motif enrichment analysis of DARs, AC-footprints did not show similar pattern of enrichment for these factors. Majority of the observed footprints overlapping genetic variants had corresponding changes between their motif and footprint scores, but the number of these instances per TF were too low for meaningful TF-specific statistical analysis. Because DARs in general were less accessible than non-DARs and their accessibility correlated strongly with footprint scores, this could be due to the lack of sensitivity when detecting footprints in DARs, resulting in the low number of observations for AC-footprints. In addition, as recently shown by a study focusing on TCF7L2 function in the hepatocytes, TF binding can be very specific even within a cell type [43]. Such highly cell type specific binding, which is still reflected on chromatin accessibility, is less likely to present with strong signals for footprint analysis or even DAR analysis with bulk ATAC-seq where the signals from several different cell types combine, which could be solved using single cell approaches [18].

In summary, differences in cell type composition, both genetically driven and not, seem to affect the identification of genetically determined TF binding from bulk ATAC-seq datasets. Scores for altered motifs overlapping footprints in DARs correspond with changes in footprint scores, which presents a strong case for the mediation by pioneer factors on the genetically determined chromatin accessibility.

## Materials and methods

### Mouse experiments

Mice were acquired from Jackson Laboratories (Boston, USA), housed at the Lab Animal Centre of the University of Eastern Finland, and experimented on under a permit from the local Animal Experiment Board (ESAVI-2015-002081). Male C57BL/6J and 129S1/SvImJ mice were fed either a chow (Teklad 2016, Envigo) or HFD (TD.88137, Harlan) from 9 weeks of age for 8 weeks. At 17 weeks of age, mice were euthanized by CO_2_ for harvesting tissues. The epididymal white adipose tissue were weighted and snap-frozen in liquid N_2_ and stored in -80°C.

### RNA extraction

Total RNA was extracted from mouse eWAT tissue using the miRNeasy Mini Kit and QIAzol Lysis Reagent from QIAGEN, according to the manufacturer’s protocol. The RNA was treated with RNase-Free DNase (QIAGEN) to remove any DNA. The RNA quality was assessed with an Agilent Bioanalyzer and the RNA 6000 Nano kit, with all samples showing a RIN score of 7 or higher.

### Library preparation and RNA-sequencing

RNA-seq libraries were generated using 800 ng of total RNA. Ribosomal RNA was first removed with the NEBNext rRNA Depletion Kit (New England BioLabs). The libraries were then prepared using the NEBNext Ultra II Directional RNA Library Prep Kit for Illumina (New England BioLabs), according to the manufacturer’s guidelines. The yield of the libraries was assessed using the Qubit DNA High Sensitivity assay (Invitrogen), and quality control was conducted with the Agilent Bioanalyzer using the DNA 1000 kit (Agilent). Finally, the indexed libraries were pooled and sequenced on the NextSeq 500 platform (Illumina) with 75 bp single-end reads.

### Nuclei extraction, library preparation and sequencing for ATAC-seq

Approximately 100 mg of eWAT tissue was used to generate libraries by following the Omni-ATAC protocol [44] with slight modifications. Nuclei were extracted using a Dounce homogenizer and 1× homogenization buffer composed of 320 mM sucrose, 0.1 mM EDTA, 0.1% NP-40, 5 mM CaCl₂, 3 mM Mg(Ac)₂, 10 mM Tris pH 7.8, 1× protease inhibitors (Roche, cOmplete), and 1 mM β-mercaptoethanol. After homogenization, the samples were filtered through a 100 µm nylon mesh filter, and nuclei were recovered by OptiPrep/iodixanol (Sigma-Aldrich) density gradient centrifugation. For the transposition reaction, 50,000 nuclei were resuspended in a transposition mix containing 25 µl 2× TD buffer, 2.5 µl transposase, 16.5 µl PBS, 0.5 µl 1% digitonin, 0.5 µl 10% Tween-20, and 5 µl H₂O. Following the transposition reaction, DNA was purified using the DNA Clean & Concentrator-5 Kit (Zymo Research). Ten microliters of the purified transposed DNA were then amplified for 9-11 cycles using 25 µl of 2× Ultra II Q5 Master Mix (New England BioLabs) and 1 µl each of the amplification primers (1.25 μM final concentration) [10]. To clean up the libraries and remove primer dimers and fragments larger than 1,000 bp, SPRIselect beads (Beckman Coulter) were used. The library yield was quantified with the NEBNext Library Quant Kit for Illumina (New England Biolabs), and quality was assessed using the Agilent Bioanalyzer and High Sensitivity DNA Kit (Agilent). Sequencing was performed using the Illumina NextSeq 500 platform with paired-end 75 bp reads.

### ChIPmentation-based ChIP-seq analysis of H3K27 acetylation

To perform ChIP-Seq on mouse eWAT, approximately 100 mg of frozen tissue was Dounce homogenized and fixed with 20 strokes using the loose pestle A followed by 20 strokes with the tight pestle B in Cell Lysis Buffer (5 mM PIPES pH 8, 85 mM KCl, 1% Igepal, 1× protease inhibitor) containing 0.5% formaldehyde. The mixture was then rotated for 8 minutes at room temperature. The cross-linking reaction was quenched by adding 150 mM glycine for 5 minutes at room temperature. The cross-linked samples were subsequently washed twice with ice-cold Cell Lysis Buffer supplemented with 1× protease inhibitors. The pellet containing the cross-linked chromatin was suspended in 300 µl of SDS Lysis Buffer (50 mM Tris-Cl, 0.5% SDS, 10 mM EDTA, 1× protease inhibitor) and incubated on ice for 1 hour. Chromatin was sheared using a Bioruptor Plus sonicator (Diagenode) for 35 cycles at a high setting (30s ON, 30s OFF per cycle), allowing the sonicator to cool down for 5 minutes after every 10 cycles. To neutralize SDS, Triton-X-100 was added to a final concentration of 1% along with 1× protease inhibitors. Samples were centrifuged at 13,000 rpm for 20 minutes, and the supernatant containing sheared chromatin was collected and divided into 140 µl aliquots for immunoprecipitation.

ChIP and library preparation followed the ChIPmentation protocol [45] with minor adjustments. For ChIP, 2 µg of Anti-Histone H3 (acetyl K27) antibody (Abcam, #4729) was added to 50 µl Protein G-coupled Dynabeads (Thermo Fisher Scientific) in 1 × PBS with 0.5% bovine serum albumin (BSA) and rotated at 40 rpm for 4 hours at 4°C. For the IgG control, 2 µg of Rabbit IgG (Diagenode, #C15410206) was used in a similar manner. Antibody-coated Dynabeads were washed three times with PBS containing 0.5% BSA and then mixed with 140 µl of chromatin samples in 1.5 ml tubes, followed by overnight rotation at 40 rpm at 4°C. Immunoprecipitated chromatin was washed with low-salt buffer (50 mM Tris-Cl, 150 mM NaCl, 0.1% SDS, 0.1% sodium deoxycholate, 1% Triton X-100, and 1 mM EDTA), high-salt buffer (50 mM Tris-Cl, 500 mM NaCl, 0.1% SDS, 0.1% sodium deoxycholate, 1% Triton X-100, and 1 mM EDTA) and LiCl buffer (10 mM Tris-Cl, 250 mM LiCl, 0.5% IGEPAL CA-630, 0.5% sodium deoxycholate, and 1 mM EDTA), followed by two washes with TE buffer (10 mM Tris-Cl and 1 mM EDTA) and two washes with ice-cold Tris-Cl pH 8.Immunoprecipitated bead-bound chromatin was then resuspended in 30 µl of 2 × TD buffer and 1 µl of transposase (Nextera, Illumina) for tagmentation. After incubation at 37°C for 10 minutes, the samples were washed twice with low-salt buffer. Bead-bound tagmented chromatin was diluted in 23 µl of water and combined with 25 µl of 2 × Ultra II Q5 Master Mix (New England BioLabs, M0544S) and 1 µl of both amplification primers [10]. Library preparation included adapter extension at 72°C for 5 minutes, reverse cross-linking at 95°C for 5 minutes, followed by 11 cycles of PCR amplification (98°C 10s, 63°C 30s, 72°C 3 minutes). After PCR amplification, double-sided purification was performed using SPRIselect beads (Beckman Coulter). The yield of the libraries was quantified using the Qubit DNA High Sensitivity assay (Invitrogen), and quality assessment was conducted using the Agilent Bioanalyzer with the DNA 1000 kit (Agilent). ChIP-seq libraries were sequenced on the HiSeq3000 (Illumina) platform with single-end 75 bp reads.

### RNA-, ATAC- and ChIP-seq preprocessing

Pseudogenome and transcriptome for the 129 mice were produced as described previously by us except for updating the Gencode transcriptome to M25 [6]. Similarly, the sequencing file preprocessing followed the described protocol apart from using STAR version 2.6.1 for aligning the reads [46].

### Statistical analysis and visualization

Statistical analyses were performed with R software (v4.1.0) [47]. Heatmaps were plotted using ComplexHeatmap (v2.13.3) and EnrichedHeatmap (v1.27.2) [48,49]. Venn diagrams were plotted using nVennR (v0.2.3) [50]. Footprint aggregate plots were generated using TOBIAS (v0.14.0) ‘plotAggregates’ function with default settings [17]. Rest of the plots were generated using ggplot2 (v3.5.1) [51].

### Outlier detection and statistical analysis of ATAC- and RNA-seq

RNA-seq and ATAC-seq samples were tested for outliers using rrcov (v1.7-5) function ‘PcaGrid’ with ‘crit.pca.distances = 0.999’ [52]. Optimal k-value for outlier detection was selected with PCAtools (v2.6.0) function ‘parallelPCA’ with ‘threshold = 0.01, max.rank = 24’ parameters [53]. Before outlier detection, for RNA-seq the read counts were normalized with VST normalization (DESeq2, v1.34.0) and for ATAC-seq with local regression fit using csaw (v1.28.0) [54,55]. After outlier removal, UMAP (umap, v0.2.10.0) was used to determine the existence of clusters deviating from the rest of the samples [56]. Differentially accessible regions (DARs) were detected using csaw with cluster as a covariate from NFRs with log_2_(CPM) > –3 counts. To focus on the Tn5 cut-sites, reads were shortened to 1bp tags at 5′-ends before read counting. For RNA-seq, DEGs were detected using DESeq2. Only genes that had at least one read in one of the samples were included in the analysis. Single cell data set with normalized counts and cell type annotation for mouse eWAT was acquired from the study by Sárvári *et al.* [20]. Cell type specific markers were detected using Seurat (v4.1.0) ‘FindAllMarkers’ command with ‘only.pos=TRUE’ [57]. Enrichment of cluster DEGs to cell type specific genes was done using clusterProfiler’s (4.2.2) enricher function. For DARs chipenrich’s (v2.17.0) ‘chipenrich’ was used for the enrichment analysis with following parameters ‘method = “ polyenrich”, genome = “ mm10”, locusdef = “ nearest_tss”’ [58,59].

### ATAC-seq analysis

Enrichment of DARs to PANTHER GO slim (v17.0) terms belonging to “ biological_process” ontology was analysed with rGREAT (v2.1.11) [60,61]. Motif enrichment analysis for the active and non-active NFRs was done using GimmeMotif’s (v0.18.0) ‘gimme motifs’ and for the different classes of DARs with ‘gimme maelstrom’ [62]. Motifs classified as “Direct” from the CIS-BP v2.0 collection and mm10 (B6) genome were used in the analyses [36]. Footprint analysis was done with TOBIAS using the strain-specific genomes [17]. For footprint analysis, the motifs were subset to those of expressed TF genes and merged with universalmotif (v1.12.3) ‘merge_similar’ with default parameters [63]. Before merging, motifs with average information content ≤0.75 were discarded to increase the reliability of the merging. Footprints from the 129 data were converted to B6 coordinates to create a consensus set of TF occupancy sites using g2gtools (v0.29) ‘convert’ [64]. Scores for B6 and 129 mice were then calculated strain specifically for these locations. Genetically altered binding sites were identified using motifbreakR (v2.8.0) with ‘filterp = TRUE, threshold = 1e-4, method = ‘ic’, legacy.score = FALSE’ and MGPv5 SNPs and InDels [65].

### RNA-seq analysis

To select active genes, zFPKM values for genes were calculated using the zFPKM R package (v1.16.0), considering a gene expressed when it had an average zFPKM > -3 and/or FDR<0.05 in one of the differential gene expression comparisons [66]. Active transcription start sites (TSSs) were detected using proActiv (v1.4.0) and the most active TSS for each gene were used in the downstream analysis as the gene start point [67]. TSSs classified as Major and Minor by proActiv were prioritized in the selection over internal promoters with higher activity score. Over-representation analysis was performed using clusterProfiler’s ‘enrichr’ using PANTHER GO slim terms (v17.0) [60].

### Integrative analysis of ATAC- and RNA-seq

NFRs were linked to the active TSS’s of genes within 1MB using ChIPpeakAnno (v3.28.0) ‘annoPeaks’ with ‘bindingRegion=c(−1e6,1e6), select=”all”’ parameters [68]. Peaks were assigned to genomic features using ChIPseeker (v1.30.3) ‘annotatePeak’ and expressed genes as annotation [69]. Correlation analyses were done using Spearman rank correlation with logCPM normalized counts from edgeR (v3.36.0) [70].

## Acknowledgements

Analyses were performed on the high-performance computing cluster provided by UEF Bioinformatics Center, University of Eastern Finland, Finland. Sequencing was partly performed using the facilities of the Genome Center of the University of Eastern Finland.

---

**S1 Fig.Batch correction resolves cell type bias from ATAC-seq data. A)** UMAP plot based on normalized DAR counts. Cluster 2 is highlighted with red circle. **B)** Chipenrich results for Cluster 2 vs Cluster 1 DARs to cell type-specific genes. Significantly enriched/depleted (FDR<0.01) gene sets labelled. **C)** UMAP plot of limma batch corrected normalized counts. **D)** Scatter plots of mean normalized DAR counts in Cluster 1 (x-axis) and Cluster 2 (y-axis) before and after batch correction.

**S2 Fig.Active NFRs present higher accessibility than non-active NFRs. A)** Boxplots of normalized counts for active and non-active NFRs in different NFR classes^1^ and B) different genomic features. Wilcoxon test P-value: * < 0.05, ** < 0.01. ^1^Common=”DAR in both diet comparisons”, HFD=”DAR in HFD comparison”, Chow=”DAR in chow comparison”, Non-DAR=“Non-DAR in both comparisons”.

**S3 Fig.Batch-corrected gene expression corresponds to chromatin accessibility at the promoter. A)** UMAP plot of normalized counts before and after batch correction. Cluster 2 highlighted with a red circle. **B)** Over representation analysis results of genes up-regulated in Cluster 2 vs Cluster 1 in sets of cell type specific genes. Bars coloured grey are non-significant (FDR>0.01). **C)** MA plots of Cluster 2 vs Cluster 1 normalized counts before and after batch correction. **D)** Circulograms for the fractions of closest NFR distance and class for expressed genes^1^. DAR=Chow-/HFD-/Common-DAR. **E)** Enrichment heatmap of ATAC-seq and H3K27ac ChIP-seq signals ±2kb from the active TSS of expressed genes. **F)** Scatter plot of normalized expression levels of genes^1^ and their distance to nearest NFR. ^1^Non-DEG=non-DEG in all comparisons, Diet=DEG only in diet comparison, Diet+strain=DEG in both diet and strain comparisons, Strain DEGs=DEG in both strain comparisons.

**S4 Fig.HFD DARs present adipocyte specific enrichment. A)** Motif enrichment for group-wise active and non-active NFRs. Heatmap columns and rows are clustered by Euclidean distance and columns are split using *K*-means clustering. Values shown are for the most significant motif for a given TF. Motifs for heatmap were selected by group-wise filtering of redundant motifs. **B)** Fisher’s test results for enrichment of Active and non-active DARs* to cell type specific genes. *Common=”DAR in both diet comparisons”, HFD=”DAR in HFD comparison”, Chow=”DAR in chow comparison”.

**S5 Fig.Footprints in liver show similar patterns as in adipose tissue. A)** Venn diagram of footprints observed in the different groups in eWAT. Numeric IDs for overlapping groups are presented in the brackets. 1=Chow B6, 2=HFD B6, 3=Chow 129 and 4=HFD 129. **B)** Scatter plot of footprint scores and normalized (log2CPM) ATAC-seq counts in the different NFRs^1^. Correlation testing with Spearman’s correlation. ^1^Common=”DAR in both diet comparisons”, HFD=”DAR in HFD comparison”, Chow=”DAR in chow comparison”, Non-DAR=“Non-DAR in both comparisons”.

**S6 Fig.Motifs enriched in DARs or non-DARs share similarity. A)** Motifs clustered by Pearson correlation. Motifs enriched in DARs in red and non-DARs in cyan. **B)** Percentages of enriched TFs in DEG-classes^1^. Counts and percentages labeled. **C)** Proportions of enriched TFs that are positive cell type markers derived from a single cell reference (Adjusted P-value<0.01 and average log2 fold change>0.25). Counts and percentages labelled. ^1^Non-DEG=non-DEG in all comparisons, Diet=DEG only in diet comparison, Diet+strain=DEG in both diet and strain comparisons, Strain DEGs=DEG in both strain comparisons.

**S7 Fig.Footprints in liver show similar patterns as in adipose tissue. A)** Venn diagram of footprints observed in the different NFRs^1^ in liver. Numeric IDs for overlapping groups are presented in the brackets. 1=Chow B6, 2=HFD B6, 3=Chow 129 and 4=HFD 129. **B)** Fractions of active and non-active NFRs by NFR-class^1^ that overlap with at least one footprint in liver. **C)** Counts for top 5 most common footprints in active and non-active NFRs^1^ in liver. **D)** Scatter plot of footprint occupancy in DARs vs non-DARs. Size of points is determined by Chi-squared P-value; red: P-value<0.05. Top 10 TFs with highest difference in occupancy fraction between DAR-class^1^ and non-DAR are labelled (P-value < 0.05). ^1^Common=”DAR in both diet comparisons”, HFD=”DAR in HFD comparison”, Chow=”DAR in chow comparison”, Non-DAR=“Non-DAR in both comparisons”

**S8 Fig.Potential genetically determined binding affects known pioneer factors in eWAT and liver. A)** Differences in eWAT footprint scores (Y-axis) across motif allelic score difference (129 vs B6) categories (X-axis). **B-C)** Word clouds of TFs with one or more synonymous footprints in NFR-classes^1^ of **B)** eWAT and **C)** liver. **D)** Violin plots for the number of AC-footprints that were detected in NFR-classes^1^ in eWAT and liver. **E-F)** Enrichment P-values (Chi-squared) of AC-footprints in NFR-classes^1^ for **E)** liver and **F)** eWAT, comparing Chow, HFD and Common DARs to Non-DARs, and Non-DARs to DARs of any class. Only TFs with P-value<0.25 shown. ^1^Common=”DAR in both diet comparisons”, HFD=”DAR in HFD comparison”, Chow=”DAR in chow comparison”, Non-DAR=“Non-DAR in both comparisons”

**S1 Table. DEGs detected in individual comparisons.** A dedicated sheet for each of the tested comparisons presented in the document. Results for all tested genes are included.

**S2 Table. Genetically altered TF binding motifs at corresponding differentially accessible footprint in liver and eWAT.** Footprint score differences presented as maximum differences between the groups included in the NFR class. Common=”DAR in both diet comparisons”, HFD=”DAR in HFD comparison”, Chow=”DAR in chow comparison”, Non-DAR=“Non-DAR in both comparisons”

